# Loss of *DYRK1A* ortholog *mbk-1* impairs locomotor behavior in *Caenorhabditis elegans*

**DOI:** 10.1101/2025.11.04.686526

**Authors:** Elysabeth Otte, Emma Bratch, Alexandria Robbins, Randall J. Roper, Charles R. Goodlett

## Abstract

Dual-specificity tyrosine phosphorylation-regulated kinase 1A (*DYRK1A*) is a dosage sensitive gene located on human chromosome 21 (Hsa21) that contributes to phenotypes associated with developmental disorders like Down syndrome and *DYRK1A* haploinsufficiency syndrome. Complete genetic knockout of *Dyrk1a* from conception is embryonic lethal, presenting a barrier in its study. The *mbk-1* gene in *Caenorhabditis elegans* has been identified as an ortholog to mammalian *Dyrk1a*, and genetic knockout of *mbk-1* in *C. elegans* is not lethal. We hypothesized that deletion of the *mbk-1* gene would alter chemosensory function, learning, and motility in *C. elegans*, and that these phenotypes would be recovered using a humanized *DYRK1A* replacement at the endogenous *mbk-1* locus. Using behavioral preference index assays, analyses of locomotion, and learning in classical conditioning procedures, an *mbk-1* knockout strain of *C. elegans*, EK228, was characterized to identify potential behavioral roles of *mbk-1*. Preference index assays assessing chemosensory capabilities determined that *mbk-1* deletion yielded no detrimental effects. Thrashing and foraging behavior analyses uncovered significant deficits in movement in the EK228 *C. elegans*, which were not present in two *mbk-1* replacement strains containing humanized *DYRK1A*, suggesting an essential role of *mbk-1* in locomotion and motility. Lastly, classical conditioning revealed no significant deficits in the abilities of the EK228 strain in forming associative connections between stimuli. Overall, these results imply functional conservation of the mbk-1/DYRK family kinases, and provide support for the use of humanized replacement strains of *C. elegans* for the study of mammalian genes.

**Graphical Abstract:** 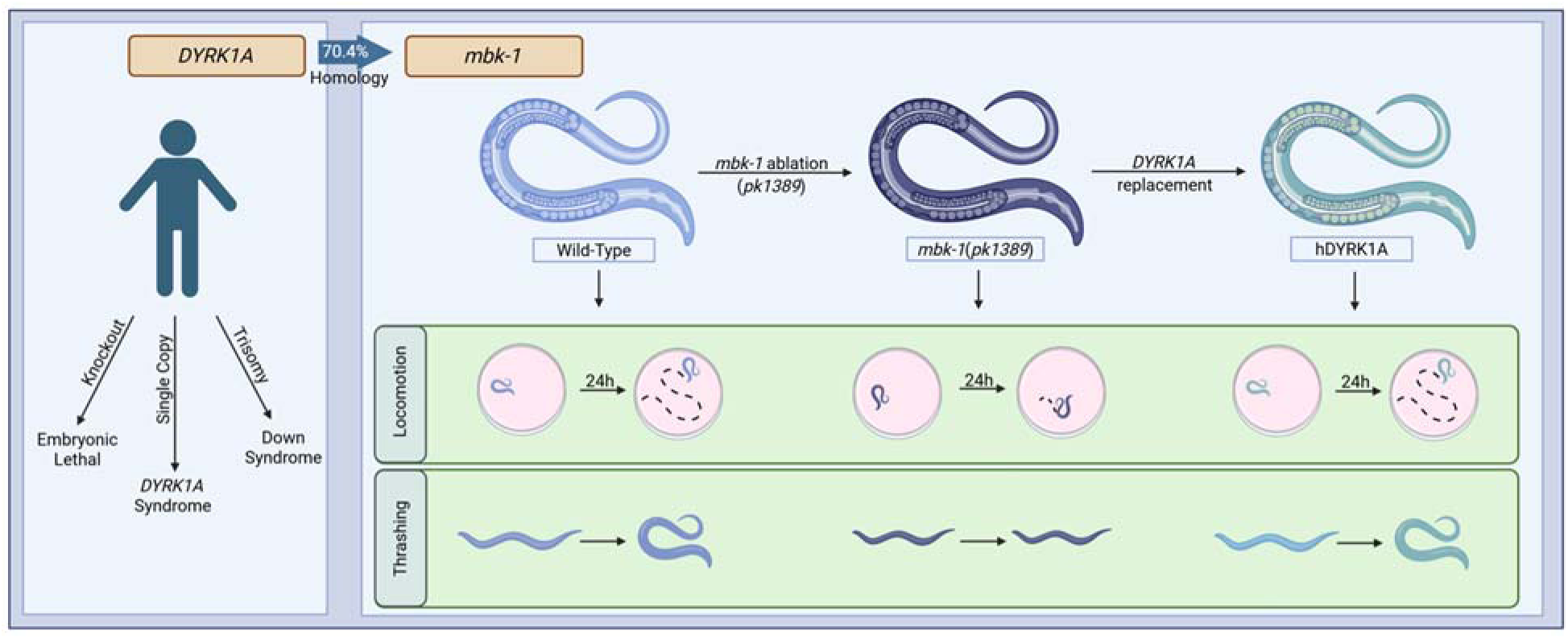
Graphical Abstract (Created with BioRender)

**Article summary:** This work provides insight into evolutionarily conserved functions of the protein kinase *DYRK1A,* which is linked to developmental disorders including Down syndrome and *DYRK1A* syndrome. *Caenorhabditis elegans* with a genetic ablation of the *DYRK1A* ortholog, *mbk-1*, were used to examine potential roles in movement, chemosensing, and associative learning and memory, and demonstrated selective deficits in thrashing and foraging locomotion that were not observed in strains with *mbk-1* replacement with humanized *DYRK1A* at the endogenous loci. The findings indicate a role of *mbk-1* in *C. elegans* in adaptive movement, which may provide insight into some of the cellular and neural mechanisms being influenced by *DYRK1A*. Understanding of the protein kinase *DYRK1A* may help to elucidate novel therapeutic pathways for developmental disorders.

## Introduction

Dual-specificity tyrosine-regulated kinase 1A (DYRK1A) is a dosage-sensitive protein that acts both to phosphorylate serine and threonine residues on other proteins and to autophosphorylate tyrosine residues in its activation loop to promote its own catalytic activity (ALVAREZ *et al*. 2007). DYRK1A is an important regulator of mammalian nervous system development, with roles in dendritic spine formation, cellular proliferation and differentiation, and in several key developmental signaling pathways (HAWLEY 2023; YANG *et al*. 2023; COURRAUD *et al*. 2025). Dysregulated DYRK1A has been found to play a key role both in the cognitive deficits associated with Down syndrome (DS) and with *DYRK1A* haploinsufficiency syndrome (*DYRK1A* syndrome), and studies have suggested a *Dyrk1a* gene-dosage-dependent effect on cognition and brain development (DANG *et al*. 2018; HAWLEY 2023). *DYRK1A* is one of approximately 225 genes triplicated on human chromosome 21 (Hsa21) in DS, and its overexpression is believed to play a role in the development of cognitive phenotypes associated with DS (GUPTA *et al*. 2016; ANTONARAKIS *et al*. 2020). In contrast, underexpression of the gene in *DYRK1A* syndrome, usually caused by mutation, results in cognitive impairment, epilepsy, and microcephaly (VAN BON *et al*. 2015). Using humans and mammalian models, studies have examined the effects of both monosomy and trisomy of *DYRK1A* and other Hsa21 genes on cognition and brain development (VAN BON *et al*. 2015; HAWLEY 2023). However, one major limitation exists in using only mammalian models to study this gene. Complete knockout and depletion of *Dyrk1a* from conception is embryonic lethal in mammalian models (DIERSSEN AND DE LAGRAN 2006), presenting barriers for determining the mechanisms that underlie the critical role of *Dyrk1a* in mammalian processes. The monetary and temporal constraints of producing and using mammalian models combined with this limited utility provide a high incentive to utilize non-mammalian model organisms that generate quickly and have simpler anatomy to study essential functions of the *DYRK1A* gene.

The minibrain (*mnb*) gene was first discovered in *Drosophila melanogaster* after an extensive search for mutants with altered brain structure (TEJEDOR *et al*. 1995). The discovery of *mnb* led to identification and initial characterizations of the human orthologous gene, *DYRK1A*. Like mammalian *DYRK1A*, *mnb* was found to play a role in dendritic spine formation in *D. melanogaster* brains, with reduction of gene expression resulting in a decreased cortical mass (ORI-MCKENNEY *et al*. 2016). Mutation of this gene results in alterations in the central brain and visual and olfactory centers, resulting in olfactory phenotypes in mutants. Additional studies in *D. melanogaster* have found that ablation of this gene results in learning and memory deficits and disrupts endocytosis processes (HEISENBERG *et al*. 1985; HELFRICH 1986; ALTAFAJ *et al*. 2001).

While *D. melanogaster* has provided an important model for studying the impacts of *mnb/Dyrk1a* on neural development and cognition, the model has limitations regarding connectome visualization and characterization. Orthologs of *DYRK1A* have been identified in all eukaryotic species examined to date, including the model organism *C. elegans* (BRODY 1997; RAICH *et al*. 2003). Located on the nematode X chromosome, minibrain-related kinase 1 (*mbk-1*) is approximately 70.4% conserved from its mammalian counterpart. As such, it has been predicted to enable serine/threonine kinase activity (WORMBASE ; RAICH *et al*. 2003). *mbk-1* is one of three *DYRK1A*/*minibrain* family members that affect the *C. elegans* nervous system, each with potentially redundant functionalities. The other two members of this family – *mbk-2* and *hpk-1* – have each been the focus of characterization studies, though complete ablation of *mbk-2* results in a 100% penetrant maternal-effect embryonic lethality. *hpk-1* null mutants display dysregulation in the expression of genes associated with aging, and have found roles of the *hpk-1* gene in stress regulation (RAICH *et al*. 2003; BERBER *et al*. 2016; LAZARO-PENA *et al*. 2023). Studies examining the effects of *mbk-1* overexpression have noted chemotaxis abnormalities, and previous studies have also established roles of *mbk-1* as a novel regulator of *C. elegans* longevity, as well as in the regulation of fatty acid desaturation and pathogen defense (MACK *et al*. 2017; MACK *et al*. 2022). Previous large-scale phenotyping efforts have included *mbk-1* ablation and loss-of-function strains among broader analyses of *C. elegans* behavior variation (RAICH *et al*. 2003; MCDIARMID *et al*. 2020). Additionally, a previous study identified several potential *C. elegans* mutants with motor-deficiency phenotypes, including one *mbk-1* ablation model containing the *ok402* mutation (RB677) (SCHMEISSER *et al*. 2017).

However, studies targeting locomotor, chemosensory, and learning phenotypes associated with loss of *mbk-1* function remain limited. Given the established role of the mammalian *mbk-1* ortholog, *DYRK1A*, in neurodevelopment and nervous system function, further characterization of *mbk-1*-dependent behavioral phenotypes may provide insight into conserved functions of the DYRK family.

The EK228 strain of *C. elegans* provides a model of *mbk-1* ablation with the *pk1389* mutation, allowing for the examination of the roles of this gene in an array of behaviors associated with nematode survival, including chemotaxis and locomotion. Humanized *C. elegans* models have emerged as a powerful tool for evaluating the functional consequences of human genetic variants *in vivo* by replacing endogenous worm genes with their human orthologs (ISLAM *et al*. 2025). In this study, we aim to expand upon the characterization of the EK228 *mbk-1* ablation model, to determine whether *mbk-1* is essential for behavior and movement, which may inform functions of the *DYRK1A* gene in mammals. We hypothesize that ablation of *mbk-1* will result in alterations in chemosensory function, movement, and learning, and confirmation of one or more of these outcomes would implicate a fundamental role for mammalian *DYRK1A*s. Additionally, we hypothesize that the replacement of endogenous *mbk-1* with humanized *DYRK1A* will improve these deficits when compared to the EK228 strain. These studies provide new evidence from this nematode model that confirm essential roles of *mbk-1* in expression of adaptive movement and further implicate the potential role of *DYRK1A* in phenotypes associated with disorders involving dysregulated *DYRK1A* including DS and *DYRK1A* syndrome. In addition, we anticipate these studies will provide behavioral models to advance understanding of potential neural and learning impacts attributed to *mbk-1*.

## Methods

### *C. elegans* strain maintenance

N2 (WBStrain00000001) and EK228 (WBStrain00007134) (Fig. 1) *C. elegans* were obtained via the *Caenorhabditis* Genetics Center (CGC) at the University of Minnesota. New samples of each strain were ordered approximately every 6 months to ensure worm health and to prevent effects of genetic drift. Humanized *DYRK1A* replacement strains were obtained from InVivo Biosystems (Eugene, OR, USA). These strains were generated through the replacement of the endogenous *C. elegans mbk-1* locus with the human *DYRK1A* coding sequence using CRISPR-Cas9-mediated genome engineering. Site-specific genome modifications were introduced using guide RNAs targeting the endogenous locus. A wild-type *DYRK1A* replacement strain (COP2302) and a *DYRK1A*(R467Q) disease-related variant strain (COP2310) were used in this study. The COP2310 R467Q strain contains a missense variant within the human *DYRK1A* sequence in the humanized locus resulting in an arginine to glutamine mutation near the C-terminal end of the kinase domain. This mutation is believed to reduce the efficiency of DYRK1A kinase binding and is commonly associated with human *DYRK1A* syndrome (EVERS *et al*. 2017). Inclusion of both the COP2302 and COP2310 strains allows us to determine if human *DYRK1A* is functionally conserved to *mbk-1*, and if variants related to human disease interfere with in vivo kinase function in the *C. elegans* model. Strain identities were validated via PCR and DNA sequencing done by InVivo Biosystems prior to distribution (see supplemental materials). The generation of these strains utilized a humanized *C. elegans* gene-replacement strategy similar to previously described approaches for functional assessment of human disease-associated variants (ISLAM *et al*. 2025). Worm strains were maintained on standard agar plates containing sodium chloride and streaked with NA22 *E. coli* as a food source in a controlled environment kept at approximately 22°C. *E. coli* stocks were re-grown every 30 days to maintain bacterial health. Worms were fed by removing a small chunk of worm covered agar from an existing plate and transferring it to a new plate with a fresh NA22 lawn.

**Figure 1.**
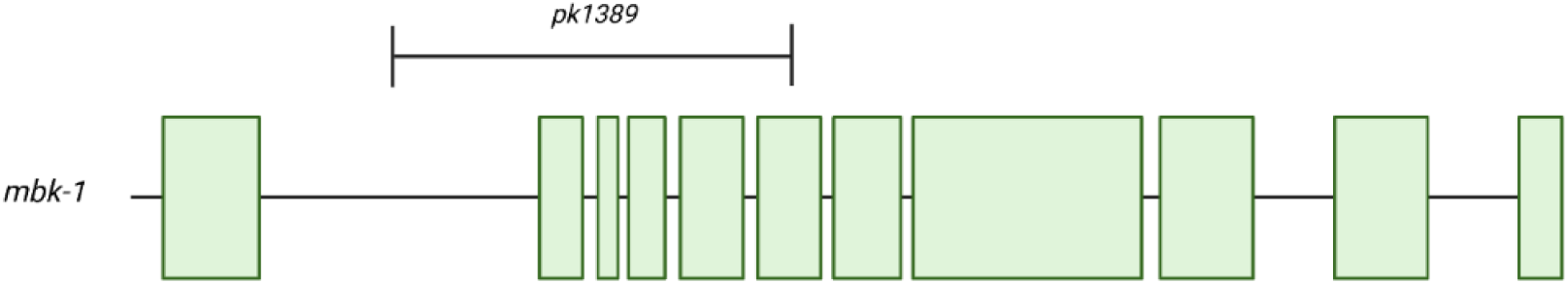
Representative schematic of the *mbk-1* gene with the *pk1389* deletion ranging from the first intron into the sixth exon, resulting in the EK228 *mbk-1* knockout strain. Created with BioRender.

### Age synchronization

*C. elegans* were washed from their maintenance plates using 15 mL sterile water, followed by centrifugation at 2500 RPM for 2 minutes. The supernatant containing NA22 *E. coli* was carefully aspirated, and the worm pellet was resuspended in 10 mL of sterile water. The tubes were inverted and centrifuged again to remove any remaining *E. coli*. The washing process was repeated once more to ensure complete removal of the bacteria.

Synchronization solution was created by combining 5 mL sodium hypochlorite, 1.25 mL 10M NaOH, and 18.75 mL sterile water. After aspirating the supernatant, 4 mL of synchronization solution was added to the worm pellet. The mixture was vigorously shaken for 5 seconds then placed on a nutating table. Transfer pipettes were used to monitor the dissolution progress every minute, starting at 5 minutes after the synchronization solution was added. Once all adult worms were dissolved, leaving only eggs in the solution, 8 mL of M9 solution were added to halt the dissolution process. The tubes were shaken and centrifuged at 2500 RPM for 2 minutes. The supernatant was aspirated, and the washing process was repeated with 10 mL M9 solution four more times.

After final wash, 10 mL M9 were added, and the tubes were placed on the nutating table for 18-24 hours to allow the eggs to hatch. Once the worms had hatched, the worms were concentrated via centrifugation and aspiration and then placed on maintenance plates with fresh NA22 *E. coli* lawns. Adolescent *C. elegans* were allowed to grow for 3 days and were tested after reaching the gravid developmental stage, just before laying their first eggs.

### Binary Choice Preference Index

A binary choice preference index assay was conducted to assess the chemosensory responses of the EK228 strain of *C. elegans* compared to the wild-type N2 strain. To prepare the assay plates, 6-well plates with wells of 35 mm diameter (Falcon 351146) were filled with a salt-free agar solution and allowed to solidify at room temperature. Each well was divided in half using a translucent template with a marked line that was taped to the bottom of the plate. On one side of each well, 4 µL of a test substance [either 0.1% benzaldehyde solution, NA22 *E. coli*, or sterile Milli-Q water (as a control)] was spotted in a 1 mm diameter target area; on the opposite side, 4 µL of sterilized Milli-Q water was spotted on 1 mm diameter target area as a neutral alternative (Fig. 2A). The substances were allowed to absorb into the agar for 2 h to establish a concentration gradient.

**Figure 2.**
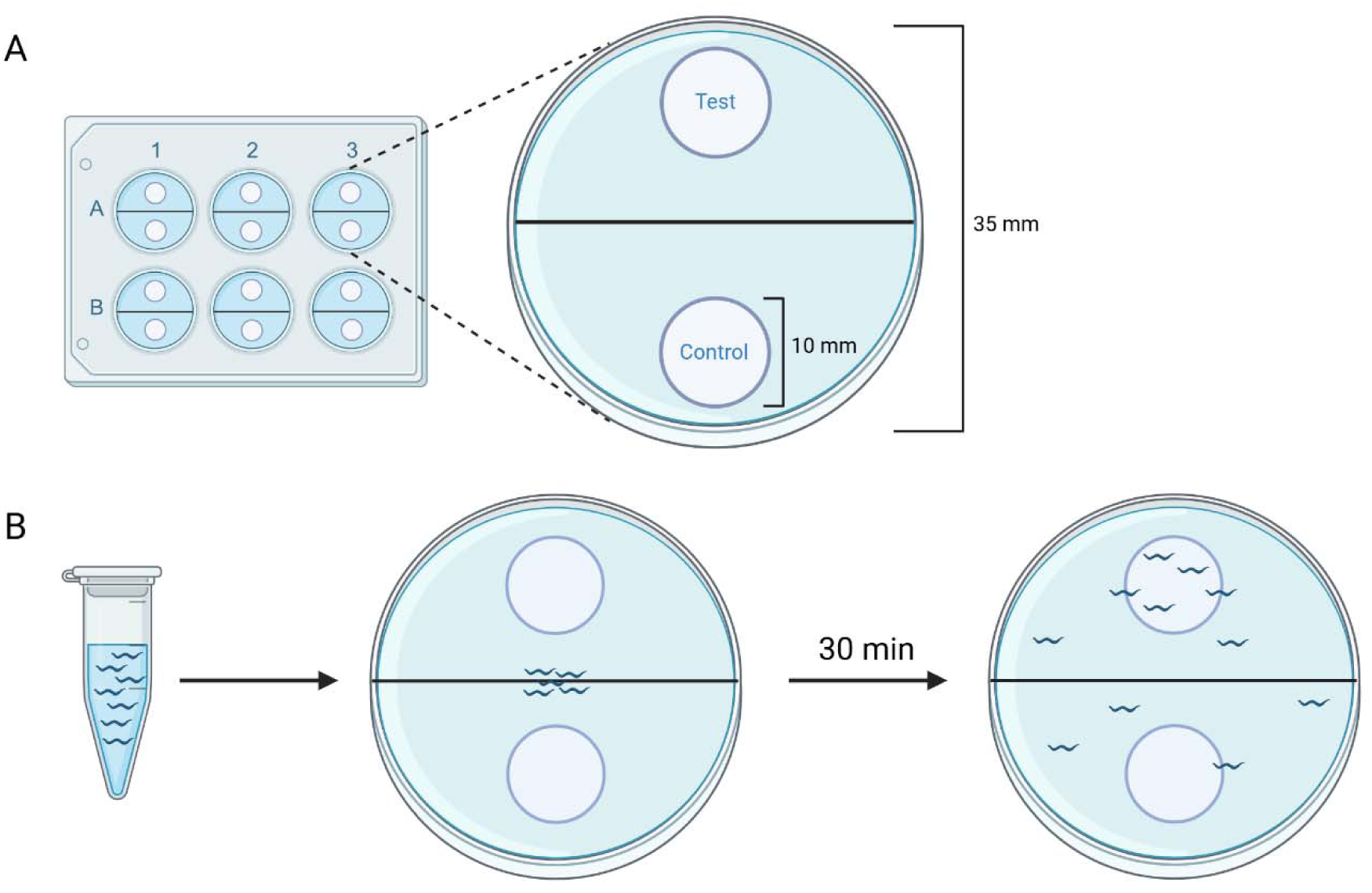
Testing setup and procedure for binary choice assay for chemotaxis. Diagram showing (A) the well setup using a standard 6-well plate and the dimensions of a well (area = 962.1 mm^2^) and the target zones (area of each = 78.5 mm^2^, 8.2% of the total); and (B) testing procedure for the binary choice assays beginning with age-synchronized worms that are washed in a 1.5 mL microcentrifuge tube, followed by plating on the test plate for the 30-minute testing period. Created with BioRender.

For the preparation of *C. elegans*, worms were washed from their maintenance plates using three aliquots of 5 mL sterile water, transferring them into 15 mL conical tubes (Crystalgen 23-2265). The tubes were then centrifuged at 2500 RPM for 2 minutes, and the supernatant was aspirated. The worms were washed two additional times with 10 mL of sterile water before being diluted to a 1:3 ratio in sterile water for the final plating.

In the assay setup (Fig. 2B), 4 µL of the cleaned worms were added to the center of each well in the 6-well plates, and any excess water was removed using a Kimwipe. The timer was started after all excess water had been removed, allowing the worms to roam freely in the wells for a duration of 30 minutes. Throughout this period, images were taken every 10 minutes using a refractive light box, and the number of worms located in each 1 cm concentration gradient zone was counted. To calculate the preference index (PI), the number of worms in the target zone was divided by the combined number of worms in both the target and control zones. In cases where a plate contained no worms in either zone, we assigned a neutral preference index score of 0.5.

### Body Bends / Thrashing

*C. elegans* were washed off maintenance plates using 15 mL of sterile water. The supernatant was then aspirated, and the worms were washed two additional times with 10 mL of sterile water. After the final wash, the worms were diluted in a 1:3 ratio, and 2 µL of the worm suspension was spotted onto a standard microscope slide. To provide the worms with space to move, an additional 2 µL of sterile water was added. The microscope slide was immediately placed under a microscope (Bausch & Lomb ASZ25L3; Nikon SMZ800), and a 20-second video of the worms’ movements was recorded at 25X magnification. The 20-second recorded video was subsequently played back at 25% speed, and the movements of the worms beyond an angle of 120 degrees were noted. The process was then repeated for the second worm strain beginning with the wash steps, and the order of strain testing alternated between trials.

The initial study of body bends used N2 (N=98) and EK228 (N=91) strains (tested in multiple replicates) and indicated a significant deficit in EK228 as compared to N2 worms (p = 0.004; **d**=0.42). To extend the study to test the hypothesis that replacement with humanized *DYRK1A* could rescue the deficit, new replications were performed with the COP2302 and COP2310 strains along with additional N2 and EK228 worms. Data across all replications were combined and the four strains were analyzed with a one-way ANOVA.

Foraging and Locomotion:

Using a titanium worm pick (Thomas Scientific, 1233S71), individual gravid adult worms were plated separately onto a lawn of NA22 *E. coli* in 10 cm petri dishes. After a 24-hour foraging period, the path of foraging movement through the *E. coli* lawn was observed, traced, and measured with Image J (SCHNEIDER *et al*. 2012). As with the body bends study, the initial foraging locomotion study with N2 and EK228 worms (N=5 worms in each group) indicated deficits in the knockout strain (p=0.001; **d**=3.14), so new replications were performed that included COP2302 and COP2310 strains together with additional N2 and EK228 worms. Data across all replications were combined and the four strains were analyzed with a one-way ANOVA.

### Classical Conditioning

#### Overview

Conditioning procedures were conducted in age-synchronized gravid adult worms using a modified version of a previously published protocol (KAUFFMAN *et al*. 2011). Five different training groups were included (Fig. 3A). Cohorts designated for classical conditioning were removed from *E. coli* food for 60 minutes, then given paired exposure in 10-cm plates of 10% butanone in 95% EtOH (the conditioned stimulus, CS) together with a restored NA22 *E. coli* lawn (the unconditioned stimulus, US) for 60 minutes. After paired exposure, cohorts given a binary preference test between 10% butanone vs. 90% EtOH vehicle at one of three retention intervals: 0 min, 30 minutes, or 60 minutes, using a 10 cm petri dish (Fig. 3B). Two comparison control groups were included. A CS-exposure group was given the 60-minute exposure to the 10% butanone CS on an *E. coli* lawn but without the preceding 60-minute NA22 *E. coli* deprivation, then assessed on the binary preference test. A naïve control group was given the binary preference test without either being deprived of NA22 *E. coli* or being exposed to the 10% butanone CS.

**Figure 3.**
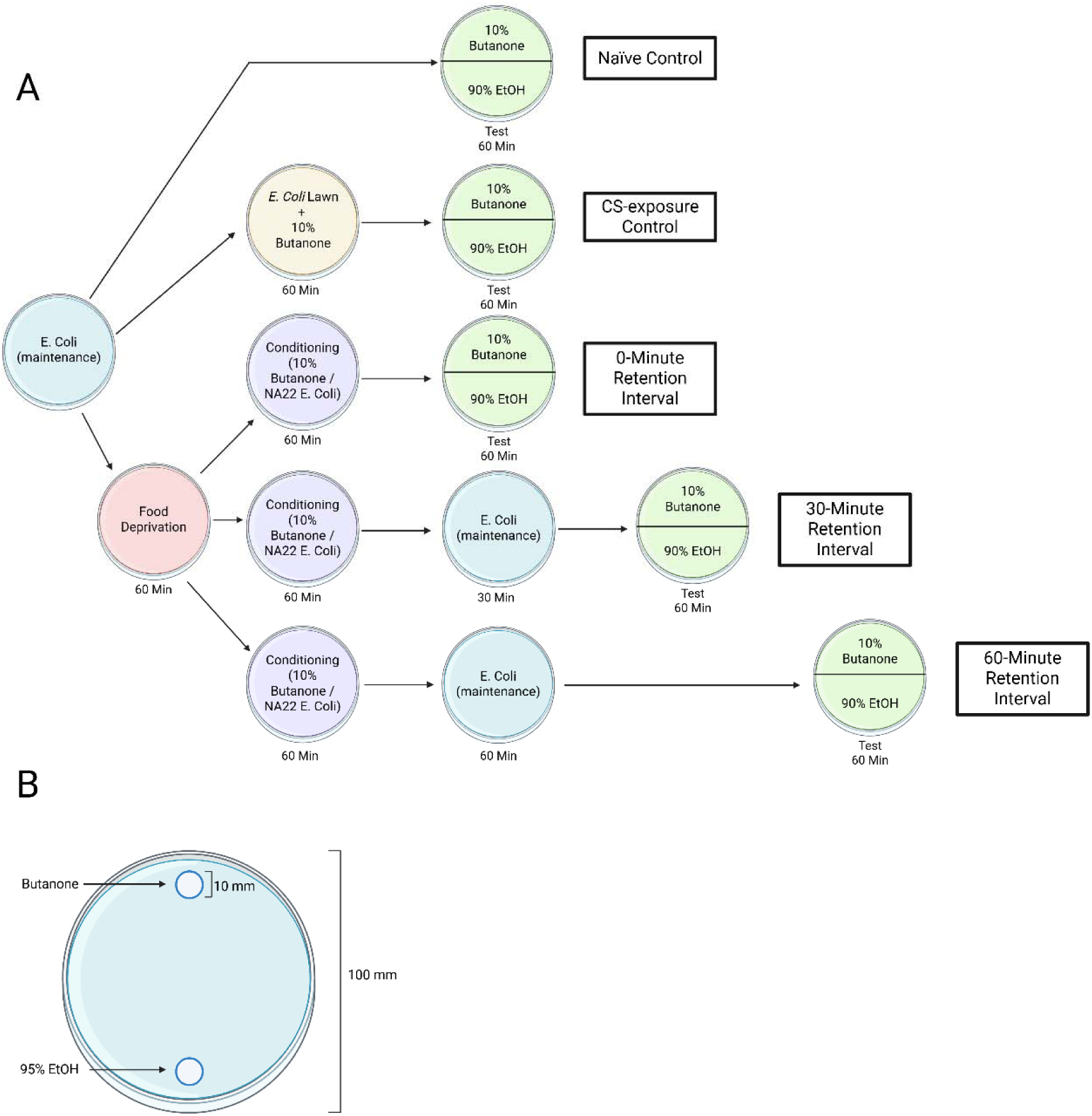
Testing setup and experimental protocol for classical associative conditioning assays. (A) Diagram of the testing plates used for conditioning assays showing the dimensions of the petri dish (area = 7854 mm^2^) and compound spotting zones (area of each = 78.5 mm^2^, 1% of the total plate area); (B) protocol for associative conditioning for the two control groups and the three conditioning groups, using butanone as CS and NA22 *E. coli* as US (B). Created with BioRender.

#### Initial collection

Worms were washed off the maintenance lawn of NA22 *E. coli* into a 15 mL conical tube using three consecutive 5 mL washes with M9 solution. After the worms settled by gravity, the supernatant was aspirated and an additional 10 mL M9 solution was added to the tubes to wash the worms. The wash cycle was repeated once more to ensure all *E. coli* were removed from the worm pellet prior to the subsequent procedures.

#### Classical conditioning procedures

After collection, the cohorts of worms designated for the CS-US paired conditioning were re-suspended in 10 mL M9 solution and placed on a nutating table for a 60-minute food deprivation period. Afterward, the worms were transferred onto 10-cm plates (200-300 uL of worms per plate) that contained bacterial lawns of NA22 *E. coli* and that had the inside of the lids streaked with 5.6 uL of 10% butanone (in 95% EtOH). This paired exposure of food-deprived worms to butanone (CS) and *E. coli* (US) extended for 60 minutes. Following the paired exposure, worms were washed off plates using 3X 5 mL M9 solution, settled by gravity, and supernatant was aspirated away. An additional 10 mL M9 solution was added to wash the worms. The wash cycle was repeated for a total of three washes in preparation for the post-conditioning test phase.

After the paired conditioning, different cohorts of worms were <u>tested for conditioned preference of 10% butanone at three different retention intervals:</u> 0 minutes, 30 minutes, and 60 minutes. The preference test for the 0-minute cohort ensued immediately following the wash cycles. Worms were diluted 1:3 using M9 solution and 15 uL of worms were plated onto 10-cm petri dishes (Fig. 3A) previously spotted with 1 uL 10% butanone in 95% EtOH at one end of the plate (target zone “A”) and with 1 uL of only 95% EtOH at the opposing end (target zone “B”).

Target zones were marked on the bottom of each plate with a dot in marker indicating the spotted zone, and the letter “A” or “B” for ease of scoring. After excess M9 was wicked away using a Kimwipe, worms were given 60 minutes to free-roam the plates. A 1% sodium azide solution (1 uL) was spotted in each target zone prior to testing, to inhibit worm movement once they entered one of the zones. After 60 minutes, images were captured of each plate, and all worms within a 1 cm diameter area surrounding the target zones were counted. For counting, each target zone was drawn on post-testing in Adobe Photoshop in order to prevent potential interference of a visible color zone in preference. For the 30-minute and 60-minute cohorts, worms were placed back onto NA22 *E. coli* maintenance lawns (in the absence of butanone) for the designated interval, then washed three times using M9 solution, diluted 1:3, and transferred onto 10 cm plates for the preference test.

Two different comparison control groups were included (Fig. 3A). The first was a naïve control group given the 60-minute preference test immediately after initial worm collection and washing. The second was a group given CS exposure in the absence of bacterial starvation.

After initial collection and washing, these worms were given a 60-minute exposure on a 10-cm plate containing an *E. coli* lawn with 5.6 uL of 10% butanone in 95% EtOH streaked onto the inside of the lid, then given the 60-minute preference test.

<u>Data analysis</u>. The preference index (PI) of 10% butanone relative to vehicle was calculated for each test plate (PI= #worms_[Target A]_ / (#worms_[TargetA]_ + #worms_[TargetB]_). PIs for butanone that are significantly greater than 0.5 (chance) and approach 1.0 in the paired groups (but not in the naïve or the CS-exposure group) would indicate behavior based on associations formed during the prior paired experience of the food-deprived *C. elegans,* i.e., learning that restored availability of bacterial food was linked to the presence of butanone, rather than resulting from inherent preferences for butanone (naïve group) or from mere exposure to the butanone CS (CS-exposure group) in the absence of bacteria deprivation.

## Results

### *mbk-1* ablation does not impair chemosensory preference at high concentrations in *C. elegans*

Previous characterization of the *mbk-1* gene suggested a role in the chemosensory abilities of *C. elegans* (RAICH *et al*. 2003). The binary choice assay of the current study quantified chemotaxis behavior toward chemo-attractive substances (relative to alternative neutral substances) over a 30-minute period by wild-type N2 and EK228 *mbk-1* knockout *C. elegans*. As shown in Fig. 4, the N2 worms displayed preference indices (PI) that were significantly greater than chance (using one-sample t-tests) for benzaldehyde and NA22 *E. coli* at 10-, 20-, and 30 minutes post-plating. For benzaldehyde, the respective means ± SEM (with p value and Cohen’s **d**) for the three time points were 0.717 ± 0.036 (p<0.001; **d**=1.06), 0.812 ± 0.034 (p<0.001; **d**=1.60), and 0.826 ± 0.039 (p<0.001; **d**=1.48). For NA22 *E. coli*, the respective values were 0.654 ± 0.048 (p=0.003; **d**=0.57), 0.692 ± 0.042 (p<0.001, **d**=0.82), and 0.796 ± 0.034 (p < 0.001, **d**=0.97). For the neutral choice test (water/water), N2 worms did not differ significantly from chance at any time point. The EK228 worms also exhibited significant preferences for the two chemo-attractants at each time point. For benzaldehyde, the PIs for the respective time points were 0.656 ± 0.046 (p=0.002; **d**=0.59), 0.765 ± 0.051 (p<.001; **d**=0.92), and 0.811 ± 0.050 (p<0.001; **d**=1.11). For NA22 *E. coli*, the respective values were 0.630 ± 0.040 (p=0.003; **d**=0.57), 0.742 ± 0.035 (p<.001; **d**=1.23), and 0.828 ± 0.027 (p<.001; **d**=2.18).

**Figure 4.**
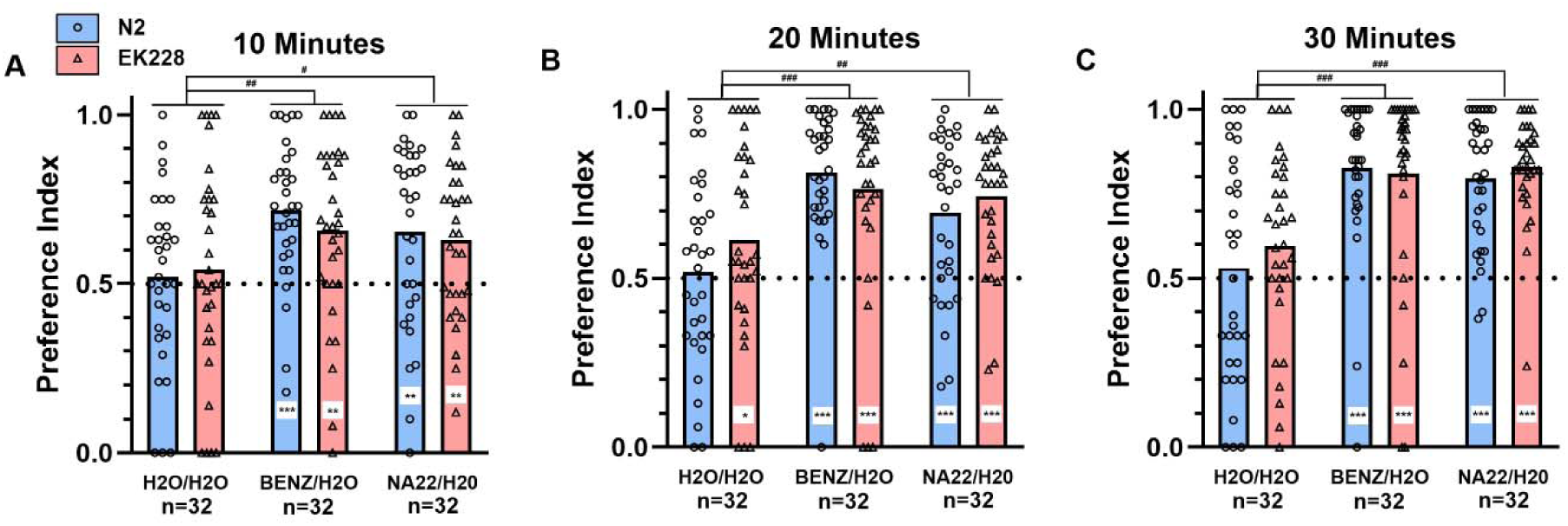
*mbk-1* mutant *C. elegans* do not display a deficit in olfaction. Preference index measurements between H_2_O and H_2_O, Benzaldehyde, or NA22 *E. coli* of N2 (N = 32 trials) and EK228 (N = 32 trials) at 10 (A), 20 (B), and 30 (C) minutes post-plating. Both strains at all three time points displayed preferences for benzaldehyde and for NA22 *E. coli,* with PIs that were significantly greater than chance (0.5) and that also were significantly greater than the PIs for groups tested on the neutral choice task (H_2_O/H_2_O). The H_2_O/H_2_O groups did not Differ significantly from chance, except for the small but significant elevation of the EK228 group at 20 minutes (PI=0.61, p<0.05). *p<0.05; **p<0.01; ***p<0.001, PI significantly greater than chance (0.5), one-sample t-test **^#^**p<0.05; **^##^**p<0.01; **^###^**p<0.001, PI significantly greater than H_2_O/H_2_O groups, Sidak post hoc comparisons of the main effect of test substances.

For the neutral choice task, EK228 worms did not differ significantly from chance at 10 or 30 minutes, but there was a small but significant difference from chance at 20 minutes [0.613 ± 0.052 (p=0.038; **d**=0.38)]. Differences in PIs between groups at each time point were analyzed with genotype × substance ANOVAs. These confirmed significant main effects of substance at each time point [10 minutes: F(2.186)=6.228, p=0.002; 20 minutes: F(2,186)=13.144, p<0.001; 30 minutes: F(2.136)=21.961, p<0.001], due to the significantly greater PIs of the benzaldehyde and NA22 choice tests as compared to the water choice test at each time point (Sidak comparisons of marginal means; see Fig. 4).

### Reductions in movement after *mbk-1* knockout are not present in strains with *mbk-1* replacement with humanized *DYRK1A*

To evaluate possible differences in thrashing movement (“body bends”) between strains, including the potential for rescue of function by replacement with humanized *DYRK1A*, twenty-second video clips of worms of the N2 (N = 141), EK228 (N = 140), COP2302 (N = 73) and COP2310 (N = 70) strains were quantified. As shown in Fig. 5A, the following average thrashing movement rates were obtained for the four strains; N2: 180.4 ± 3.0 bends per minute; EK228: 151.3 ± 3.5 bends per minute; hDYRK1A(WT) (COP2302): 168.1 ± 3.6 bends per minute; and DYRK1A(R467Q) (COP2310): 172.5 ± 3.4 bends per minute. Analysis using a one-way ANOVA and Games-Howell post hoc comparisons confirmed a significant effect of strain on body bend frequency (p < 0.001; ηp^2^ = 0.104), with the EK228 worms having significantly less thrashing than N2 (p < 0.001), COP2302 (p = 0.005), and COP2310 (p < 0.001) worms. Small but significant differences were also identified between N2 and COP2302 strains (p = 0.043). No significant differences were found in the comparison of the two humanized *DYRK1A* strains (COP2302 and COP2310; p = 0.800).

**Figure 5.**
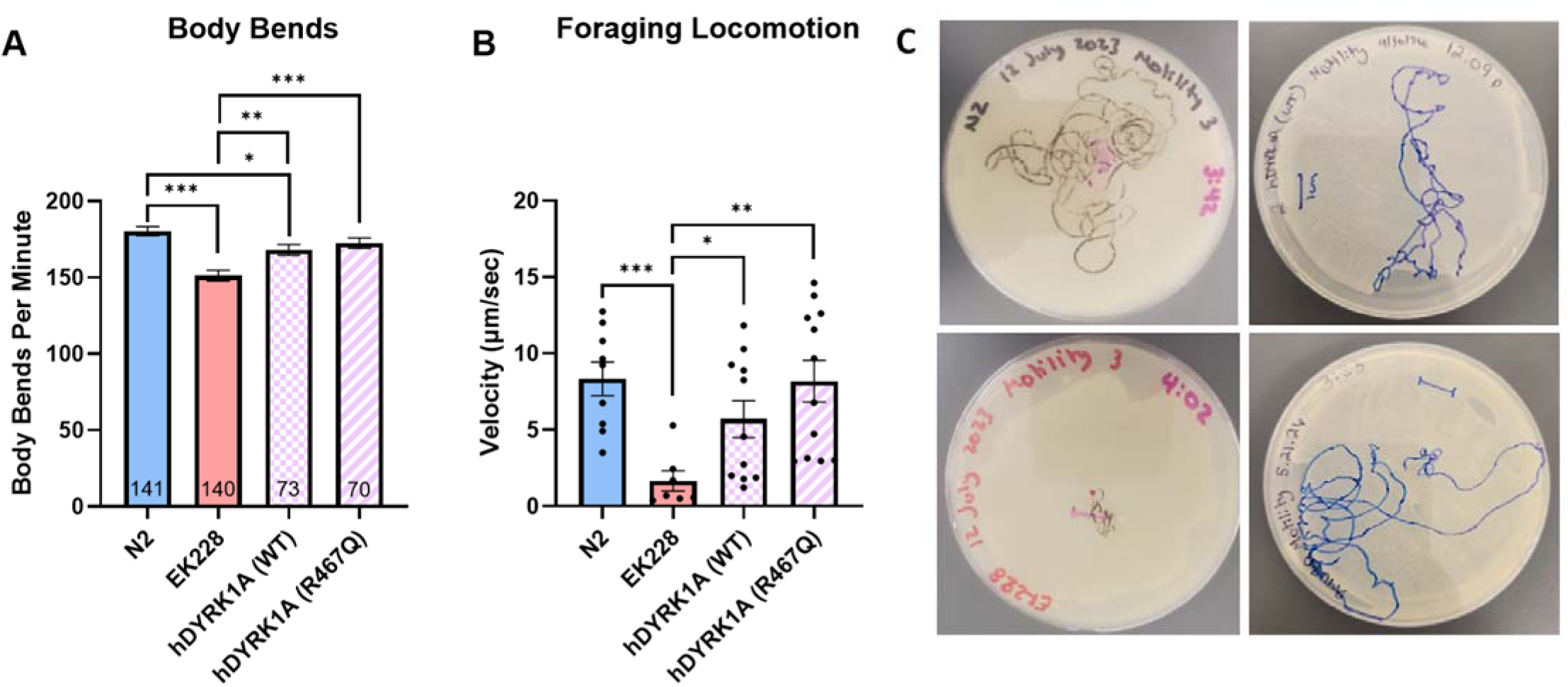
*mbk-1* ablation significantly impairs locomotion and movement, and replacement with humanized DYRK1A restores function. (A) Thrashing behavior of individual N2 (N = 141), EK228 (N = 140), hDYRK1A(WT) (COP2302) (N = 73), and hDYRK1A(R467Q) (COP2310) (N = 70) worms per minute while suspended in water (mean +/- SEM). (B) Distance traveled during forward movement of individual N2 (N = 9), EK228 (N = 7), COP2302 (N = 11), and COP2310 (N = 12) worms over 24 h in a foraging environment with access to NA22 *E. coli*, quantified as velocity (µm/sec, distance traveled over 24-hours, mean +/- SEM). (C) Representative images of movements by N2 (top left), EK228 (bottom left), COP2302 (top right), and COP2310 (bottom right) *C. elegans* over a 24-hour period. Images are of the median plate of each group. *p<0.05^, **^p<0.01, ***p<0.001, differences between averages significantly different between strains, one-way ANOVA with Games-Howell post hoc analyses

The thrashing behavior that the body bends assay assesses involves general axial movement capacity but does not correlate directly to forward movement and foraging behavior. To accomplish that, a motility assay was adapted to measure amount of movement over time in a foraging context. The average velocities of movement (µm/sec) over 24 hours on an NA22 *E. coli* lawn for N2 (N = 9) (8.34 ± 1.1 µm/sec), EK228 (N = 7) (1.65 ± 0.6 µm/sec), COP2302 (N = 11) (5.70 ± 1.2 µm/sec), and COP2310 (N = 12) (8.17 ± 1.3 µm/sec) were determined (Fig. 5B, C). One-way ANOVA analysis showed a significant main effect of strain on distance traveled over 24 hours (p = 0.003; ηp^2^ = 0.313). Games-Howell post hoc comparisons identified significant reductions in distance traveled during foraging locomotion in the EK228 strain when compared to N2 (p < 0.001), COP2302 (p = 0.047), and COP2310 worms (p = 0.003). Neither the COP2302 nor the COP2310 strains differed significantly from wild type (p = 0.398; p = 1.00, respectively), and the two humanized *DYRK1A* replacement strains did not differ significantly from each other (p = 0.538).

### Knockout of *mbk-1* has limited effects on *C. elegans* olfactory associative learning

The effect of *mbk-1* knockout in *C. elegans* on associative learning was assessed by comparing N2 and EK228 strains in acquisition and retention of classical conditioning of an olfactory preference. For the initial test of acquisition, bacteria-deprived worms were given a 60-minute paired exposure of 10% butanone (the conditioned stimulus [CS]) in the presence of the restored bacterial lawn (the unconditioned stimulus [US]). Expression of a conditioned preference for 10% butanone was assessed in a binary choice preference test immediately following conditioning. Comparison control groups included a naïve group (no manipulation) and a CS-exposure group (no bacteria deprivation). Both the N2 and the EK228 strains exhibited significant and strong conditioning for 10% butanone with PIs that were significantly above chance ([PI = 0.954; p < 0.001; **d**=12.07] and [PI = 0.898; p < 0.001; **d**=4.08], respectively), whereas the PIs of the Naïve and the CS exposure groups were not significantly above chance for either strain (Fig. 6A). A Strain × Group ANOVA yielded only a significant main effect of Group [F(2, 84)=18.61, p<0.001, η^2^p = 0.39] reflecting the higher PIs of the Paired 0-Min groups as compared to controls. Post hoc Sidak comparisons confirmed the Paired 0-minute group had significantly higher PIs than N2 Naïve or CS exposure controls. For the EK228 strain, the Paired 0-minute group had significantly greater PIs than the Naïve group, but the CS exposure group was intermediate to and not significantly different from either the Naïve or the Paired 0-minute group. The EK228 and N2 strains did not differ from each other for any of the three training conditions.

**Figure 6:**
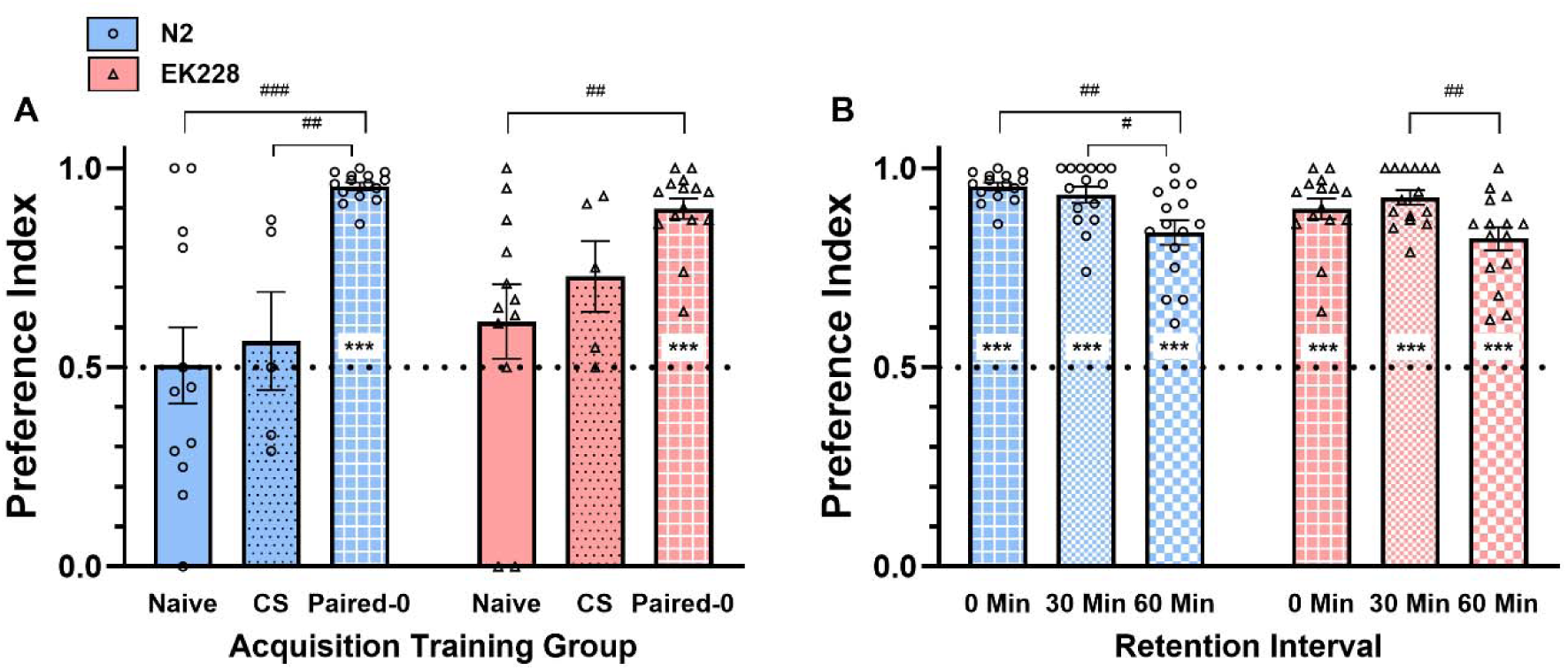
Acquisition and retention of classical conditioning of olfactory preferences. **(A)** Bacteria-deprived wild type and *mbk-1* knockout strains that were given paired exposure to 10% butanone (CS) and NA22 *E. coli* (US) then tested immediately, both showed significant and strong conditioned preference for butanone, with PIs that were significantly above chance (0.954 and 0.898, respectively). The PIs for the naïve groups or the CS exposed groups of the two strains did not differ significantly from chance. The higher PIs of the Paired-0 Min groups compared to controls yielded a significant main effect of Group [F(2, 84)=18.61, p<.001, η^2^_p_ = 0.39]. Post hoc Sidak comparisons indicated the N2 strain had significantly greater PIs for the Paired-0 group than either the Naïve or CS Exposure controls. For the EK228 strain, the PI of the Paired-0 group was significantly greater than the Naïve group, but the CS exposure group was intermediate to and not significantly different from either of the other two groups. The EK228 and N2 strains did not differ significantly from each other for any of the three training conditions. **(B)** Expression of the conditioned butanone preference declined significantly over time (main effect of interval, p<0.001). For the N2 strain, the PI at the 60-minute retention interval was significantly reduced relative to both the 0 minute and 30-minute retention interval; for the EK228 strain, the PI at 60 minutes was significantly reduced relative to 30 minutes. The N2 and EK228 strains did not differ significantly from each other at any retention interval. **\*\*\***p<0.001, one-sample t-test relative to 0.5 (chance) **^#^**p<0.05; **^##^**p<0.01; **^###^**p<0.001, Sidak post hoc comparisons

To assess whether *mbk-1* knockout could impair retention of the conditioned olfactory preference, additional groups of N2 and EK228 worms were given paired exposure, then returned to a maintenance plate for 30 or 60 minutes before testing for butanone preference. These were then compared to the 0-minute group to assess whether conditioned preferences declined with the longer retention intervals. As shown in Fig. 6B, all conditioned groups showed PIs that were significantly greater than chance at all intervals, but both strains showed a significant reduction in PIs at the 60-minute retention interval relative to earlier intervals [main effect of interval: F(2,84)=11.85, p<0.001, η^2^p = 0.22]. The N2 strain was significantly reduced relative to both 0- and 30-minute intervals, and the EK228 strain was significantly reduced relative to the 30-minute interval. EK228 and N2 strains did not differ significantly from each other at any retention interval.

## Discussion

Testing of the wild-type and *mbk-1* knockout strains revealed major locomotor deficits as a result of *mbk-1* ablation, though no deficits were identified in chemosensory preferences or in acute conditioned learning. Humanized *DYRK1A* replacement strains did not display these locomotor deficits, suggesting that localization and function of *mbk-1* in neural networks may be essential for controlling locomotion and motility necessary for foraging in the presence of bacterial food sources, but less essential for chemosensory detection or chemotaxis directed toward localized chemoattractants. Replacement of the endogenous *mbk-1* locus with human *DYRK1A* can partially restore locomotor function, though the hDYRK1A(WT) (COP2302) strain had a small but significant reduction as compared to N2 animals in the body bends assay.

Given that R467Q is a disease-associated missense variant commonly associated with *DYRK1A* syndrome and intellectual disability, this finding was unexpected. One possible explanation of these results is that the R467Q variant retains sufficient functional activity to support locomotor behavior in *C. elegans*, despite altering *DYRK1A* function in other contexts. This explanation would align with human studies reporting mild phenotypic severity associated with the R467Q mutation when compared to other *DYRK1A* missense mutations (EVERS *et al*. 2017). Additional studies examining molecular and cellular consequences of the R467Q variant will be required to determine whether this observation reflects variations in binding capacity and function.

Raich et al. first identified *mbk-1* as the closest *C. elegans* ortholog of human *DYRK1A* and demonstrated that increased *mbk-1* dosage impairs chemosensory behavior in a dose-dependent manner. These effects were observed in mature neurons and were reversible, suggesting that DYRK-family kinases can directly modulate neuronal function independent of developmental abnormalities (RAICH *et al*. 2003). Consistent with these findings, the results of this study indicate that manipulation of *mbk-1/DYRK1A* activity influences behavioral phenotypes associated with movement and foraging in adult *C. elegans* and further support an evolutionarily conserved function of DYRK-family kinases in nervous system function. This study additionally supports previous findings of impaired movement in a separate, genetically-distinct *mbk-1* ablation model (RB677) (SCHMEISSER *et al*. 2017), and extends previous findings regarding the importance of *DYRK1A* gene dosage to the functional consequences of replacing the endogenous worm kinase gene with human *DYRK1A* and a disease-associated mutant *DYRK1A* variant.

*C. elegans* locomotion and movement during foraging behavior are indicators of physiological health span, longevity, and development (ROLLINS *et al*. 2017). Previous studies have used locomotor behavior to determine nematode nervous system health (ZHANG AND CHEN 2023; PETRATOU *et al*. 2024). *DYRK1A* has been associated with neural development and behavior in humans and mammalian models. Determining potential effects of *mbk-1* mutations on the simplified nervous systems of nematodes helps to provide supportive data on the role of *DYRK1A* and may provide a basis to pursue how altered gene-dosage effects of *DYRK1A* may arise in the nervous systems of humans with *DYRK1A* syndrome or DS. The present study determined that EK228 *mbk-1* knockout worms have significantly reduced thrashing when in suspension, and a profound reduction in locomotive behaviors related to movement and foraging (Fig. 5). These effects were significantly reduced in strains carrying h*DYRK1A* gene replacement. While the exact mechanisms underlying the locomotor deficits observed in *mbk-1(pk1389) C. elegans* remain unclear, *mbk-1* has been implicated in regulation of neural development, lifespan, stress responses, and metabolic processes. Reductions in locomotion observed in the present study may reflect neuronal dysfunction, systemic physiological effects, altered energy balance, or changes in muscle function. Future studies should incorporate morphological and tissue-specific approaches to determine the relative contributions of both neural and non-neural mechanisms of *mbk-1* on locomotor function (RAICH *et al*. 2003; MACK *et al*. 2017; MACK *et al*. 2022) The results of this study indicate a deficit in locomotion and mobility during extended foraging periods, raising the possibility that similar deficits may influence any assessments requiring extended locomotor responses in other contexts, such as long-term preference testing or olfaction studies. A limitation of the present study is the potential contribution of background genetic variation to phenotypes observed in the *mbk-1(pk1389)* strain. Though background effects cannot be completely ruled out, the EK228 strain has been reported to have undergone six rounds of outcrossing, reducing the likelihood that phenotypes arise solely from unrelated background mutations. Both humanized *DYRK1A* replacement strains demonstrated significantly greater motility than the EK228 knockout strain, supporting a role for *mbk-1/DYRK1A* signaling in the regulation of locomotor behavior. These findings suggest that the observed phenotype is not solely attributable to background mutations, although future studies utilizing additional independent *mbk-1* alleles or extensively backcrossed strains would further strengthen this conclusion.

The results of this study complement previous large-scale behavioral phenotyping studies that included *mbk-1* mutant strains by providing a focused analysis of locomotion, chemotaxis, associative learning, and foraging-related behaviors. While broad phenotypic approaches are useful for identifying candidate phenotypes across genetic backgrounds, the present study allowed for the examination of specific behaviors relevant to conserved *DYRK1A* function. The observed locomotor impairments, improved with humanized *DYRK1A* strains, support a role of *mbk-1* in the regulation of locomotor behavior, while suggesting that not all sensory or associative learning processes are fully dependent on *mbk-1* function.

*C. elegans* locomotor behavior depends on integration of sensorimotor information involving both neuronal and muscle functionality. Potential explanations for the impaired movement in EK228 *C. elegans* may include defects in dopaminergic signaling, cholinergic transmission, synaptic vesicle release, muscle responsiveness, or upstream motor circuit integration. Identifying a role of *mbk-1* in one or more of these processes of neurotransmission may provide new insights into its role in neural control of adaptive movement. Exposure to a food source slows the general movement of *C. elegans*, an effect known as a basal slowing response (BSR), which is regulated by dopaminergic neurotransmission (PETRATOU *et al*. 2024). An exacerbated BSR in EK228 worms may be indicative of a role of *mbk-1* in the regulation of the dopaminergic system of nematodes. The alterations in locomotor activity and movement in *mbk-1* deficient *C. elegans* may additionally indicate an evolutionarily-conserved function of *DYRK1A* in the dopaminergic system, which plays a crucial role in regulating locomotor function in nematodes.

*D. melanogaster* strains with various *mnb* mutations display altered sense of olfaction. This is supported by the impact of *mnb* deletion on the olfactory centers of the fly brain (TEJEDOR *et al*. 1995). In *C. elegans,* chemosensing is a primitive sensory system comprised of four organs: amphid, phasmid, inner labial, and outer labial organs. Various neurons are involved in the sensing of different substances. For example, AWC neurons mediate attraction to butanone, and a combination of AWC, ASE, ADF, AWA, and BAG neurons are involved in the chemotaxis towards a food source such as *E. coli* (HART AND CHAO 2010). Previously, the EK228 strain of *C. elegans* was described to display a slight but significant defect in attraction to low doses of olfactory cues as a result of the null mutation in the *mbk-1* gene, though *mbk-1* ablation was not associated with olfactory neuron proliferation or differentiation (RAICH *et al*. 2003). In the present study, preference index determination of EK228 *mbk-1* mutants using relatively high concentration gradients of two known chemoattractants showed a lack of significant defects in chemosensory function. Both N2 and EK228 had a significantly increased preference index for target zones over chance by 10 minutes (Fig. 4). By testing both an AWC-specific, as well as a neural system that integrates multiple chemosensory pathways, the EK228 strain was determined to lack overall olfactory deficits. Future studies should utilize lower concentrations of chemoattractants in order to examine if this lack of effect persists, as well as molecular studies to determine whether dopaminergic, cholinergic, or other neuronal pathways contribute to the observed phenotype.

Altered *DYRK1A* gene dosage levels have been associated with cognitive impairment and the development of learning deficits in mammals. Our results indicate no impairments in associative learning in a classical conditioning assay were present in the EK228 strain *of C. elegans*. Previously reported phenotypes associated with drosophila *mnb* mutants found learning impairments (HEISENBERG *et al*. 1985). For this *C. elegans* model, the possibility cannot be ruled out that compensatory genes may be present (i.e., *mbk-2*) that may limit the effects of *mbk-1* ablation on phenotypes that were unaffected in this study (e.g., chemosensory responses; classical conditioning).

Similarly to benzaldehyde, the AWC olfactory neuron – specifically the AWCON neuron – is responsible for the detection of the volatile odorant butanone used in this classical conditioning assay (KODAMA AND JURADO 2007). The ability of the EK228 strain to associate the presence of butanone with a food source demonstrates a lack of olfactory deficits associated with *mbk-1* ablation (Fig. 6). *C. elegans* associative learning is a complex behavior mediated through neural mechanisms including sensory neuron plasticity and glutamatergic signaling (STETAK AND HAJNAL 2011; MCMILLEN AND CHEW 2024). EK228 worms showed no deficit in associative learning following testing, with preference indices for both N2 and EK228 strains being significantly greater than chance (0.5) (Fig. 6). These results may suggest compensatory effects of other *minibrain* family genes related to *mbk-1* in *C. elegans or* may highlight the need to test longer retention intervals (e.g., up to 24 hours) to assess consolidation into a more enduring form of long-term memory.

In summary, the findings of this study demonstrate that disruption of *mbk-1* selectively affects locomotor movement and foraging speed in *C. elegans*, while leaving chemotaxis and short-term associative learning largely unaffected under the tested conditions. Reduction of the locomotor phenotypes associated with *mbk-1* ablation by humanized *DYRK1A* replacement strains supports a conserved functional relationship between *mbk-1* and its mammalian ortholog. Although the molecular and cellular mechanisms underlying these behavioral effects remain to be determined, this study contributes to our understanding of *mbk-1* function in vivo and establishes a framework for future investigations into *DYRK1A* processes. These results support the utility of *C. elegans* as a model for examining conserved aspects of *DYRK1A* biology and for exploring the behavioral consequences of human disease-associated *DYRK1A* variants.

## Data Availability Statement

The authors affirm that all data necessary for confirming the conclusions of the article are present within the article and figures.

## Acknowledgements

We thank Daniel Omura, Dr. Bethany Neal-Beliveau, and Dr. Simon Katner for their contributions to the development and execution of this study, as well as Morgan Peters and Rachel Mack for their assistance with data collection. We thank Ben Jussila and InVivo Biosystems for assistance in the acquisition of the COP2302 and COP2310 strains, and Dr. Thomas Caulfield and Digital Ether Computing for their use of the COP2302 and COP2310 strains.

