## Supplemental Tables and Figures for "Loss of *DYRK1A* ortholog *mbk-1* impairs locomotor behavior in *Caenorhabditis elegans*"

Supplemental Materials:

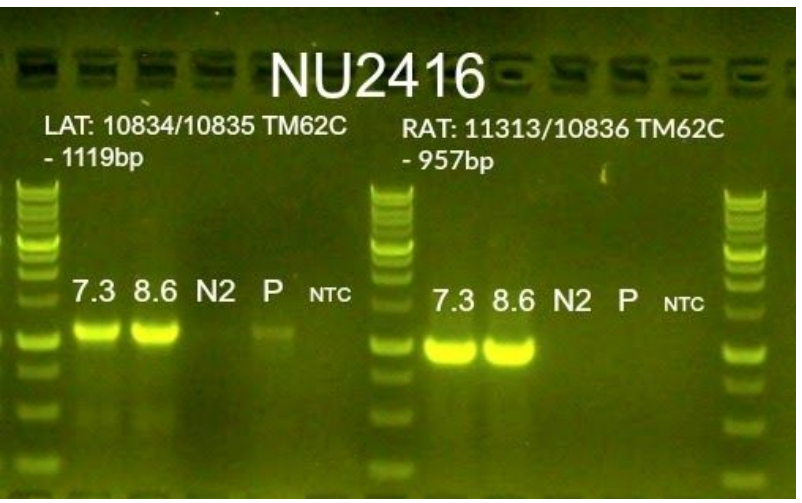

SF1: PCR validation provided by InVivo Biosystems for the hDYRK1A(WT) strain COP2302 (shown as 7.3) showing successful insertion of the *DYRK1A* gene when compared to N2 control in left (LAT) and right (RAT) homology arm tests.

| **Strain** | **Genotype** | **Validation** |
| --- | --- | --- |
| COP2302 | Humanized *DYRK1A* (WT) replacement of *mbk-1* | PCR and Sanger sequencing |
| COP2310 | Humanized *DYRK1A* replacement containing *p.R467Q* variant | HRMA screening and Sanger sequencing |

ST1: Table summarizing humanized DYRK1A strains and strain validation methods used by InVivo Biosystems prior to strain acquisition.
